## Supplementary figures for "MICA: A multi-omics method to predict gene regulatory networks in early human embryos"

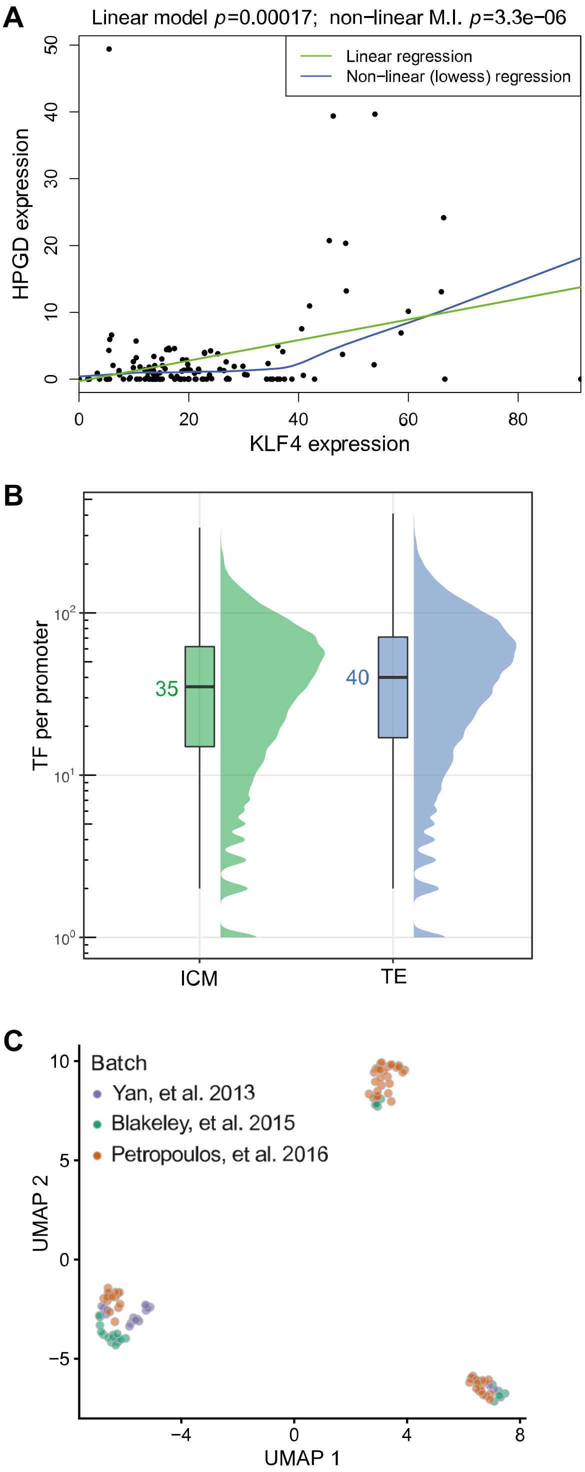


**Figure S1. (A)** Regulated genes may not ‘switch on’ until the transcription factor expression level reaches a certain threshold. This non-linear regulatory effect may be detected more efficiently using mutual information-based methods, compared to linear models. **(B)** Distribution of enriched transcription factor (TF) motifs per gene promoter. The median values are highlighted. These data were computed using the motifs enriched in footprints found within open chromatin regions in the low-input ATAC-seq from the inner cell mass (ICM) and trophectoderm (TE). **(C)** UMAP analysis of scRNA-seq data early human blastocyst stage embryos shown in Figure 3A. Here cells were coloured according to the primary data from the respective publication indicated to demonstrate that batch effects have been minimised and the scRNA-seq data is integrated irrespective of the primary study.

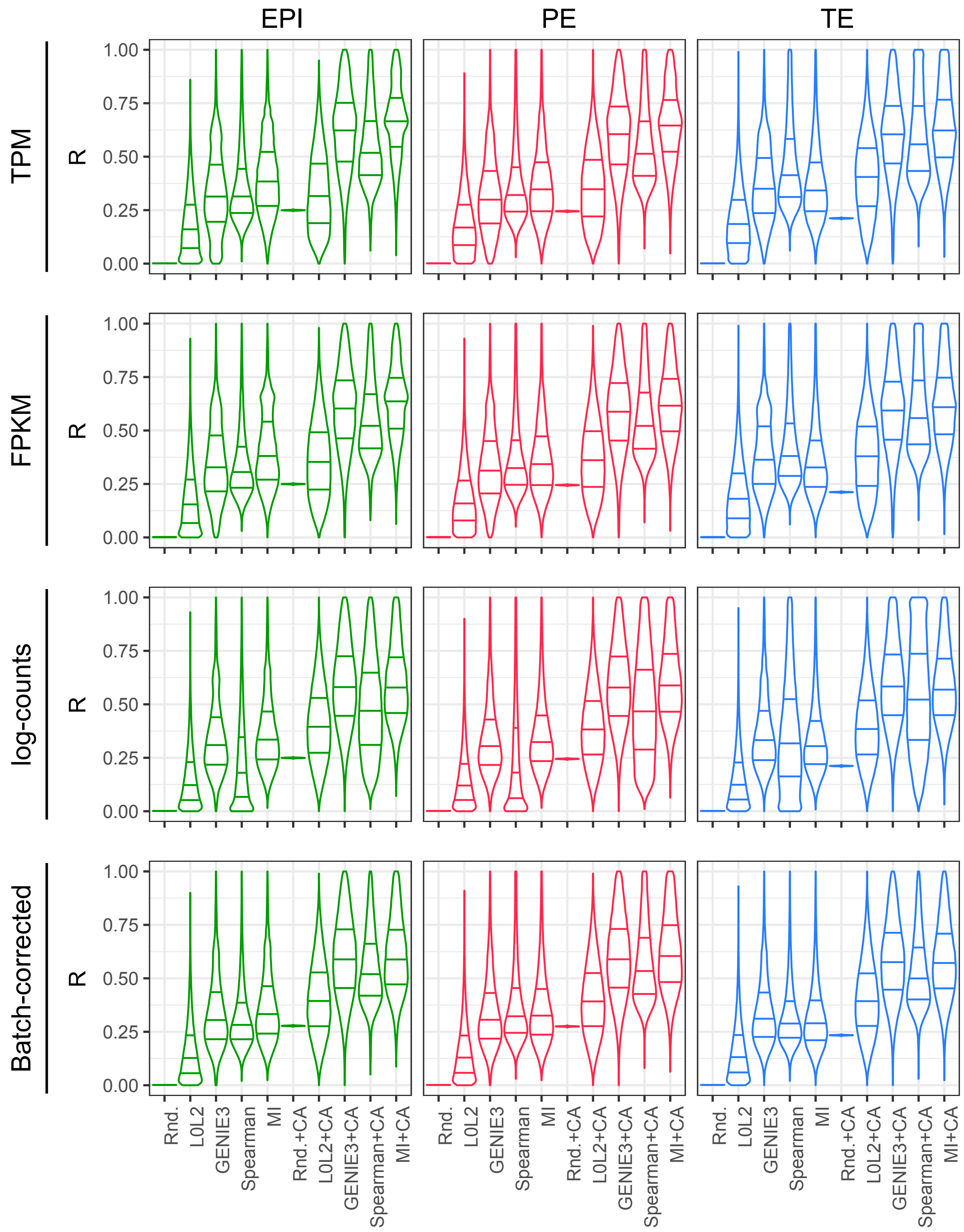


**Figure S2.** *R* distribution evaluation of the GRN prediction methods with and without chromatin accessibility (CA) refinement. The reproducibility score *R* is the bootstrap estimate of the posterior probability of observing an edge *E* given the dataset *D*, *R=P(E|D)*, for the top 100,000 predicted edges.

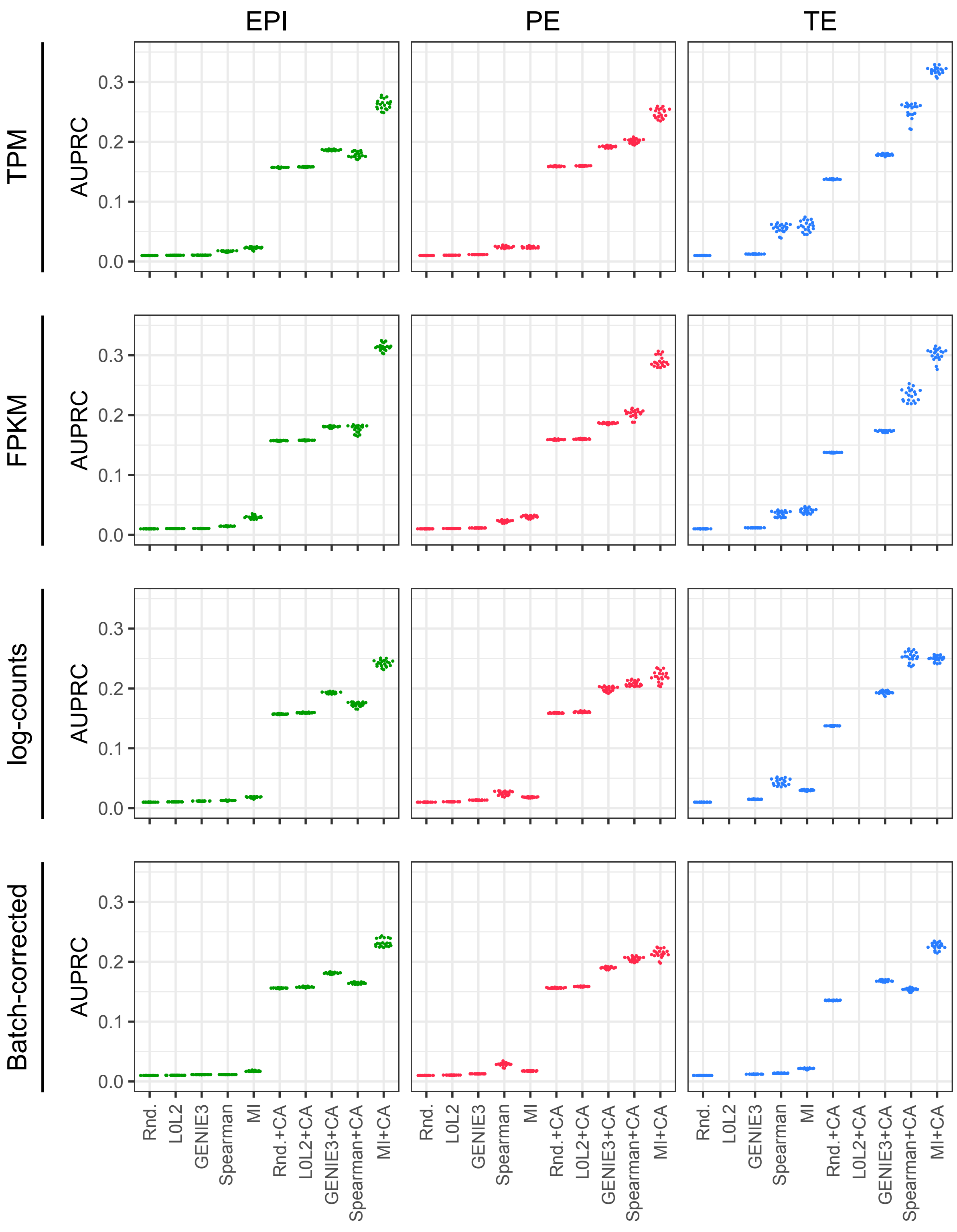


**Figure S3.** Repeated 2-fold cross-validation of the GRN prediction methods with and without chromatin accessibility (CA) refinement. The area under the precision-recall curve (AUPRC) quantifies the extent to which the interactions inferred from the first fold coincide with the top-1% interactions inferred from the second, reference fold. Cross-validation was repeated ten times. The L0L2 method didn't converge in the TE dataset.

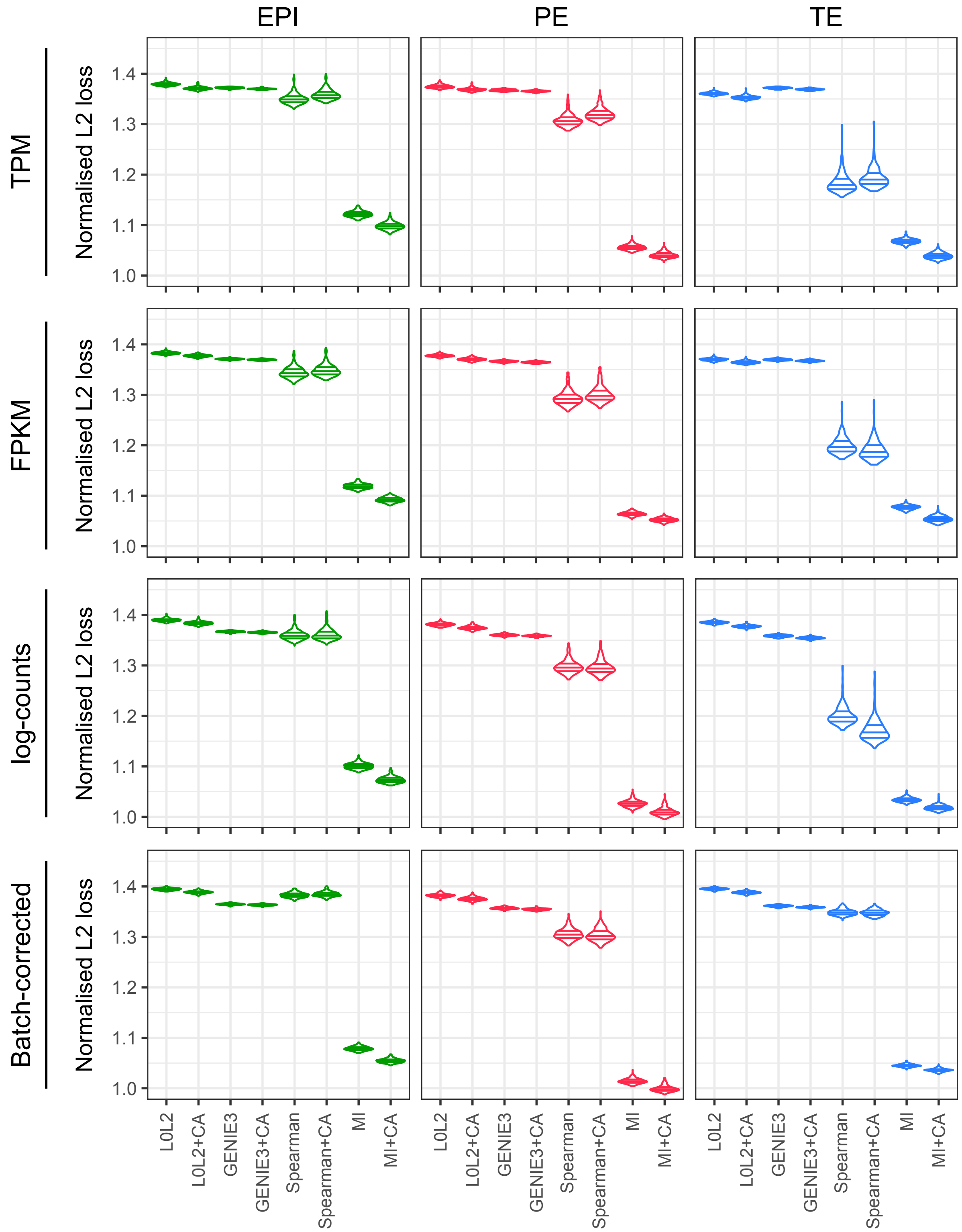


**Figure S4.** Normalised L2 loss evaluation of the GRN prediction methods with and without chromatin accessibility (CA) refinement. As in Fig. S5, 2-fold cross-validation was employed ten times, comparing the scores obtained from the application of a GRN inference method to each data split with the L2 loss function. The resulting losses were divided by the product of the square roots of the L2 norms of the network scores of each of the 2 data folds.

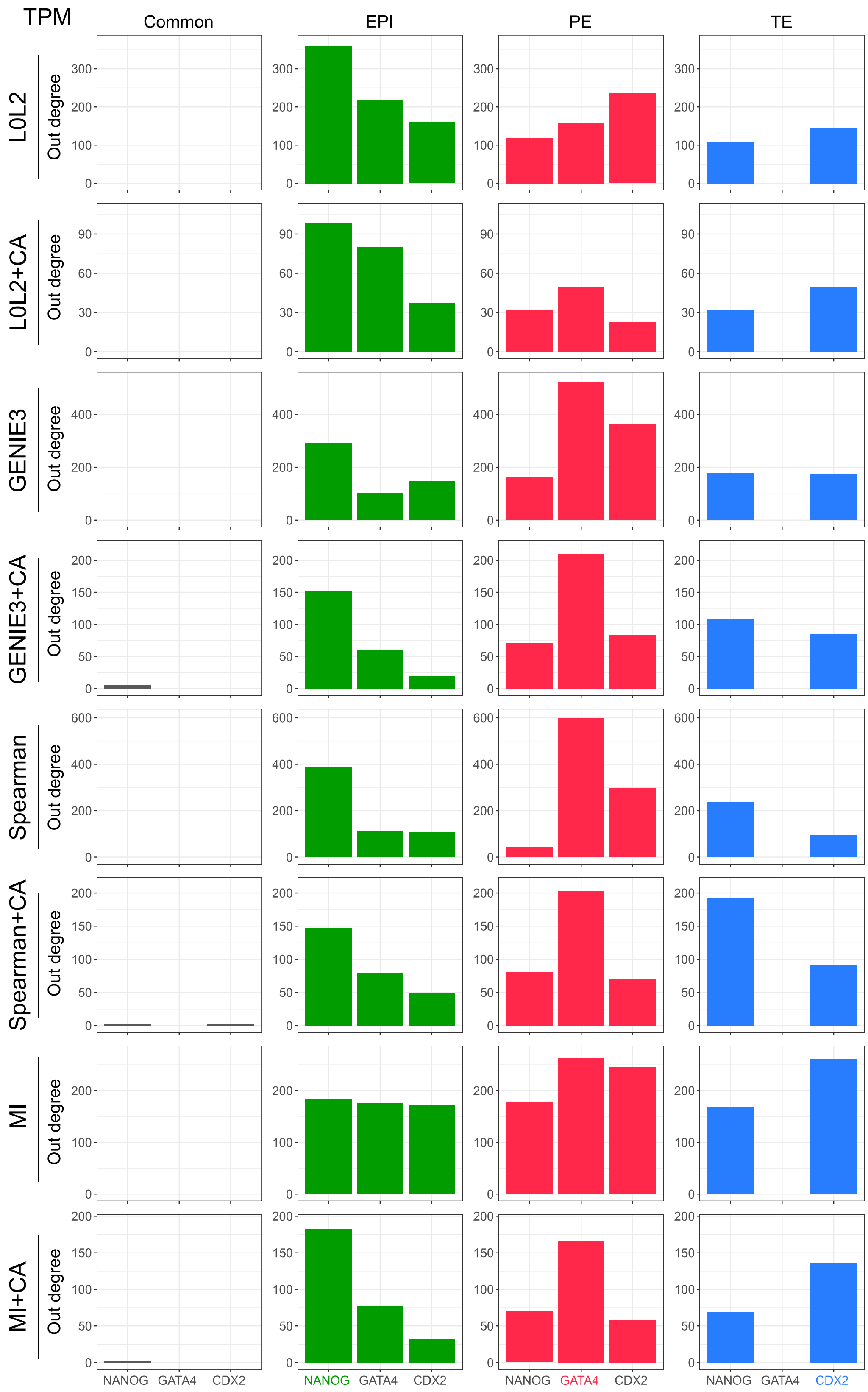


**Figure S5.** The number of genes regulated by the markers of each one of the cell types in the human blastocyst (EPI: Epiblast, PE: Primitive Endoderm and TE: Trophectoderm) is used as a proxy for marker activity in the networks predicted by the different network inference methods evaluated in this work. The results in this figure were generated with the gene expression after TPM normalisation.

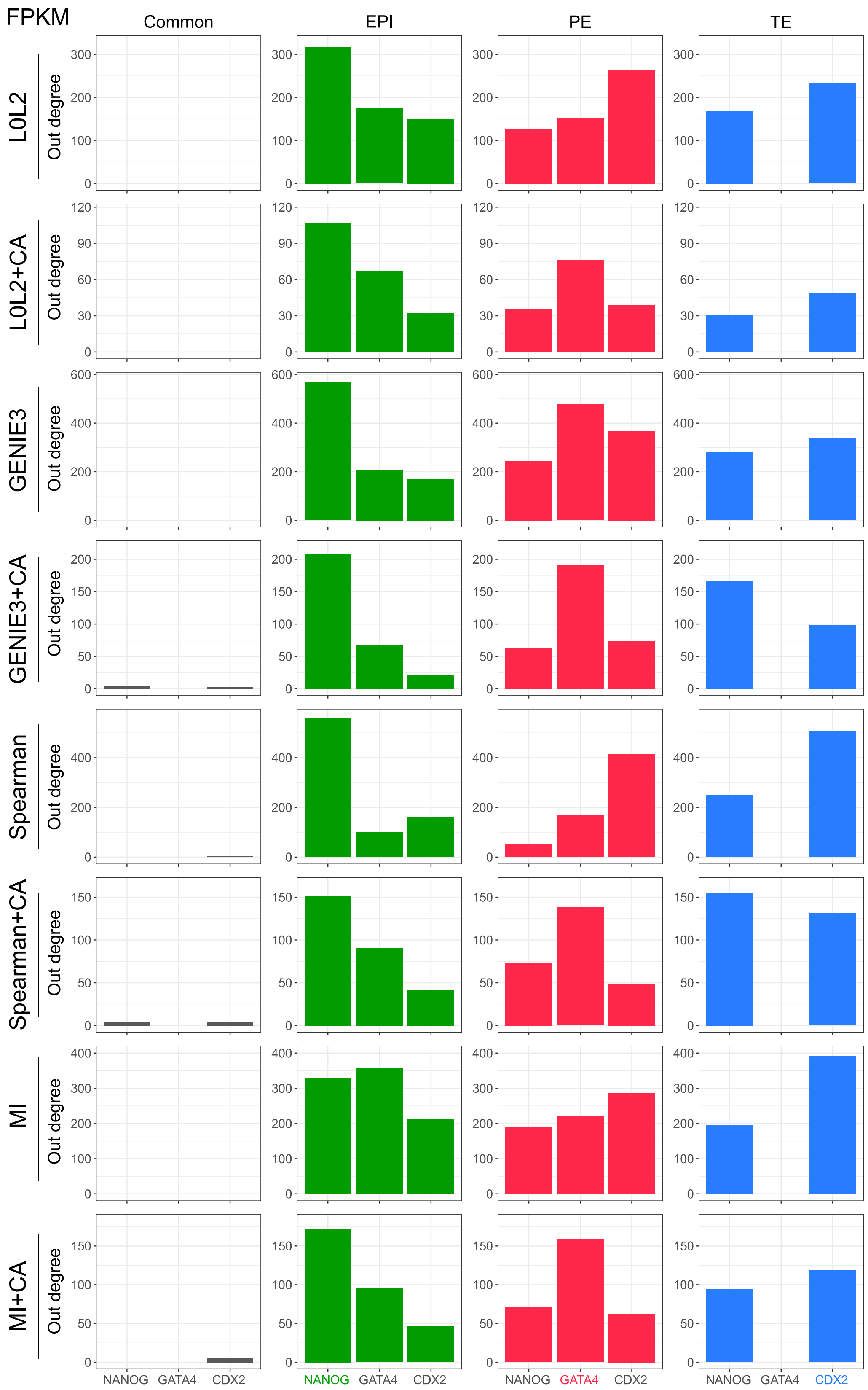


**Figure S6.** The number of genes regulated by the markers of each one of the cell types in the human blastocyst (EPI: Epiblast, PE: Primitive Endoderm and TE: Trophectoderm) is used as a proxy for marker activity in the networks predicted by the different network inference methods evaluated in this work. The results in this figure were generated with the gene expression after FPKM normalisation.

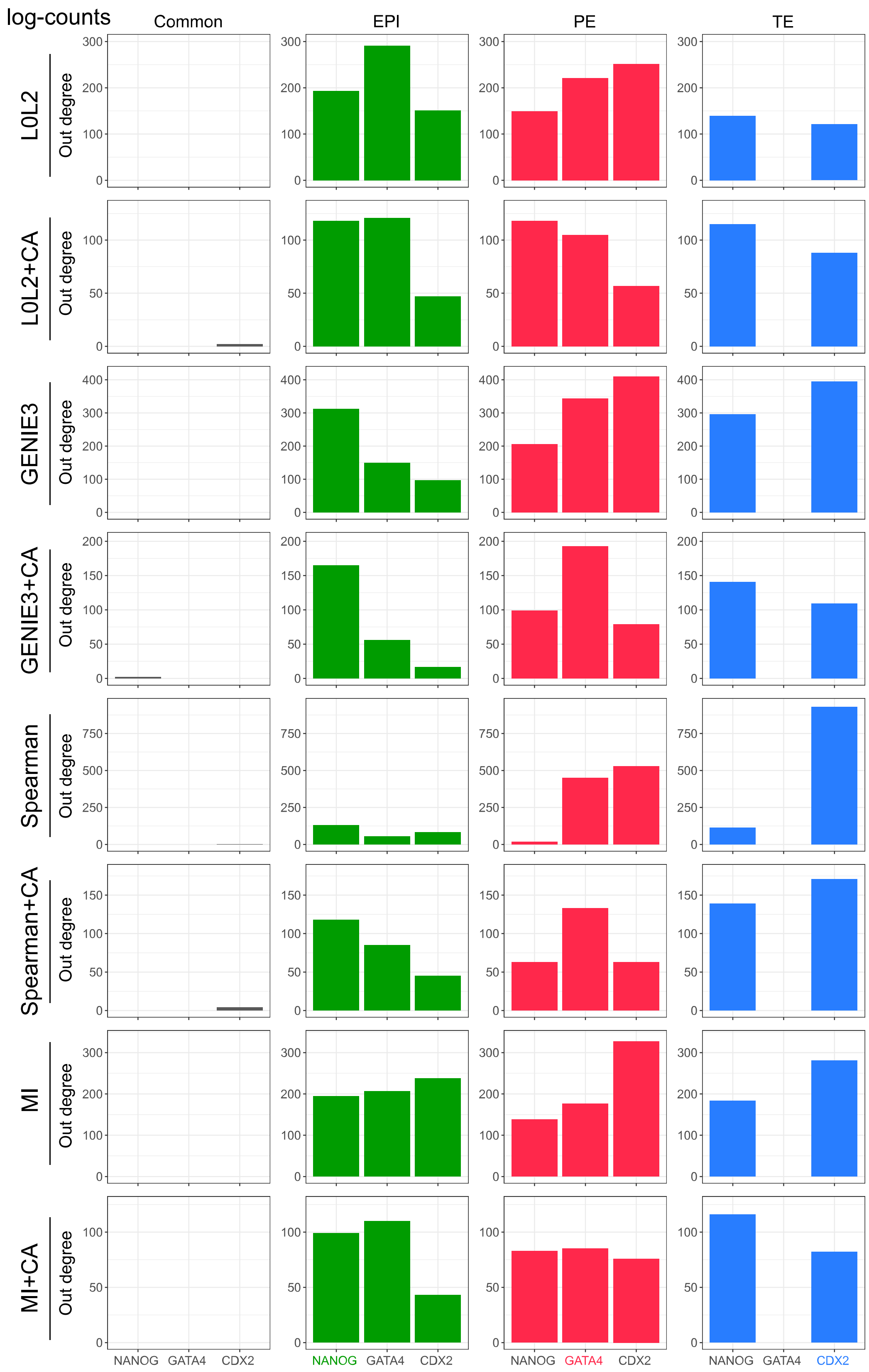


**Figure S7.** The number of genes regulated by the markers of each one of the cell types in the human blastocyst (EPI: Epiblast, PE: Primitive Endoderm and TE: Trophectoderm) is used as a proxy for marker activity in the networks predicted by the different network inference methods evaluated in this work. The results in this figure were generated with the gene expression after log-count normalisation.

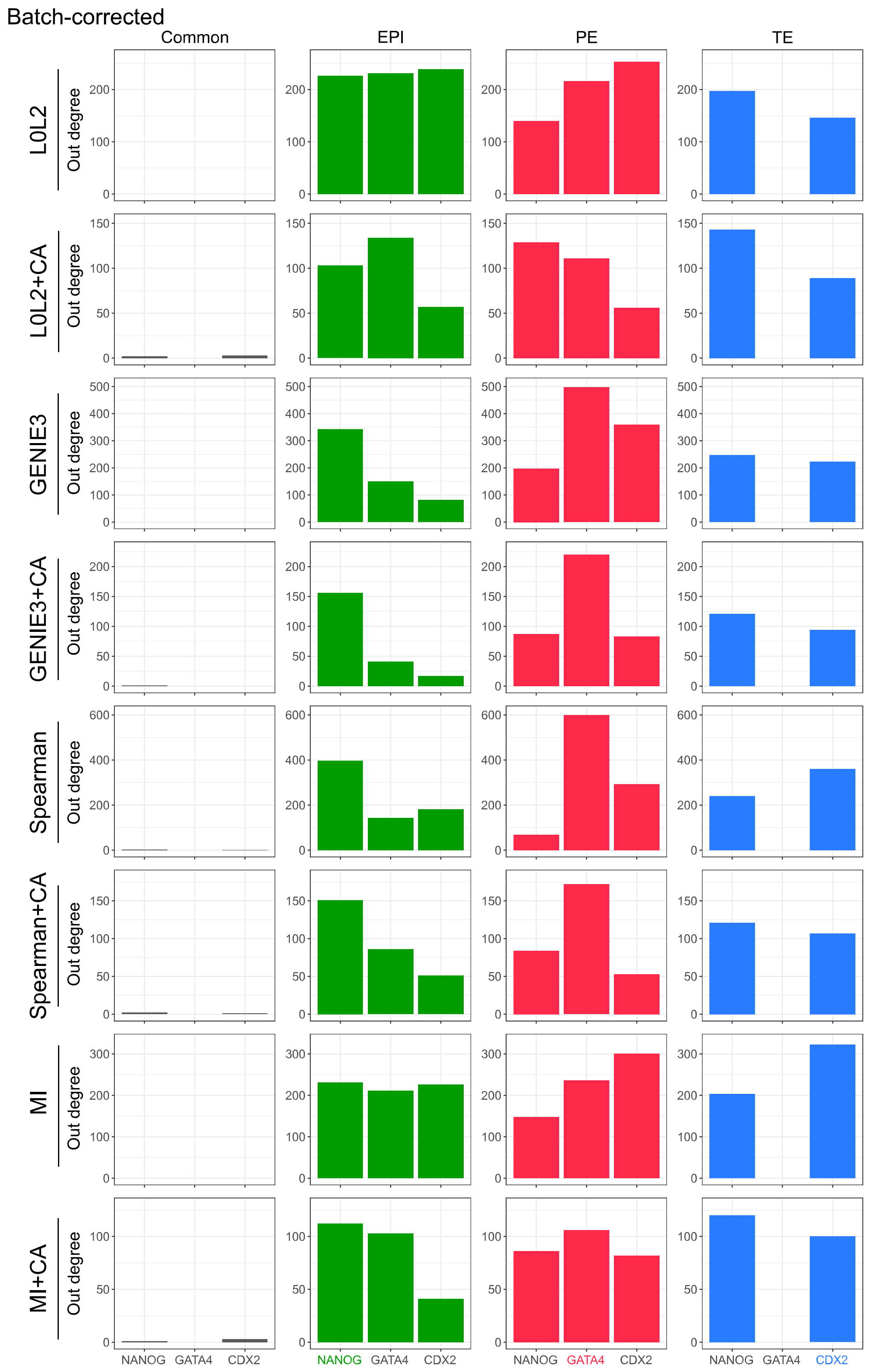


**Figure S8.** The number of genes regulated by the markers of each one of the cell types in the human blastocyst (EPI: Epiblast, PE: Primitive Endoderm and TE: Trophectoderm) is used as a proxy for marker activity in the networks predicted by the different network inference methods evaluated in this work. The results in this figure were generated with the batch-corrected gene expression values.

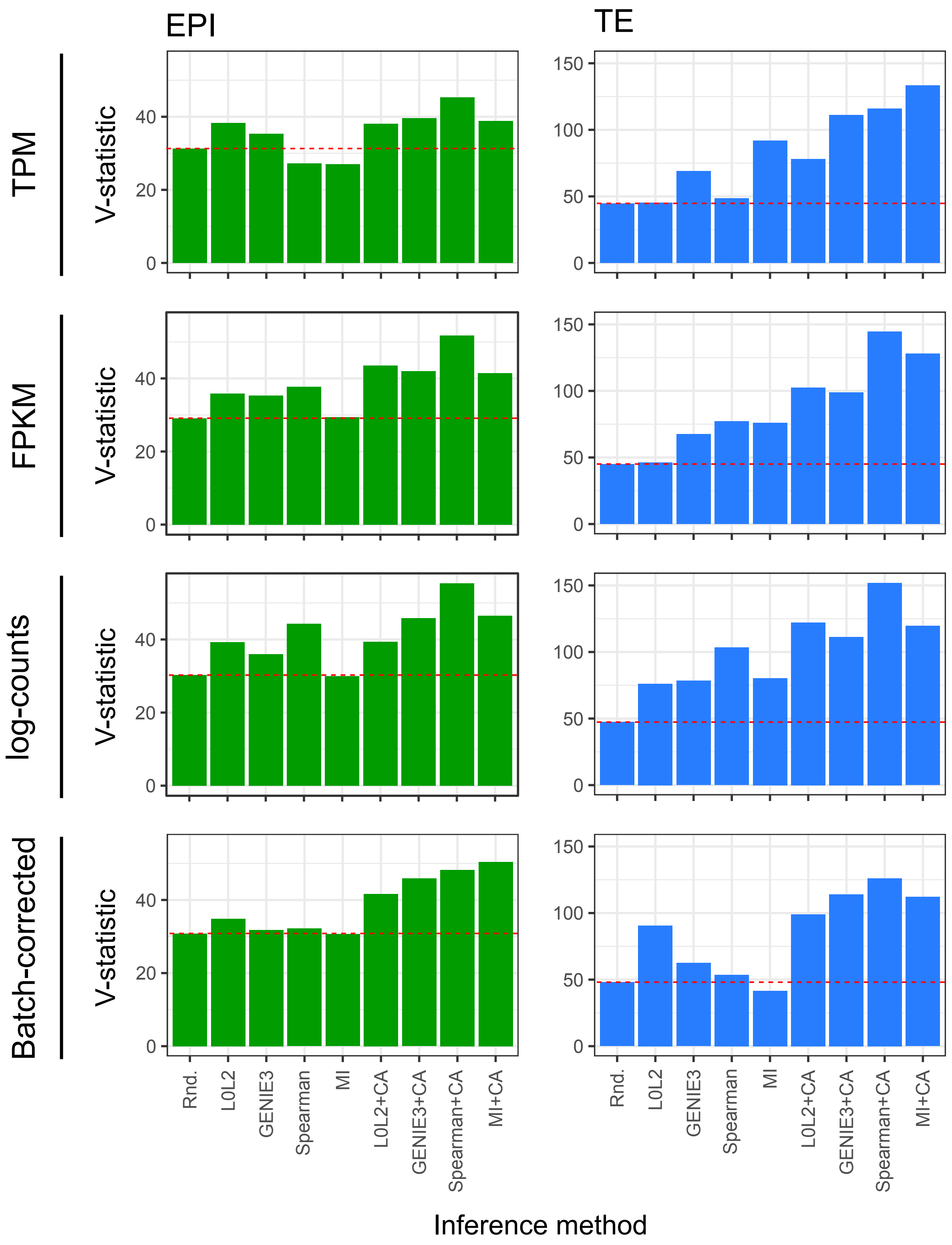


**Figure S9.** The V-statistics represent the number of relevant gene-sets that were enriched at a significance level of 10% ($p$<0.1) in a gene-set enrichment analysis performed with the genes regulated in the 25 top-predicted epiblast or trophectoderm interactions. in each of the networks predicted by the different network inference methods evaluated in this work.

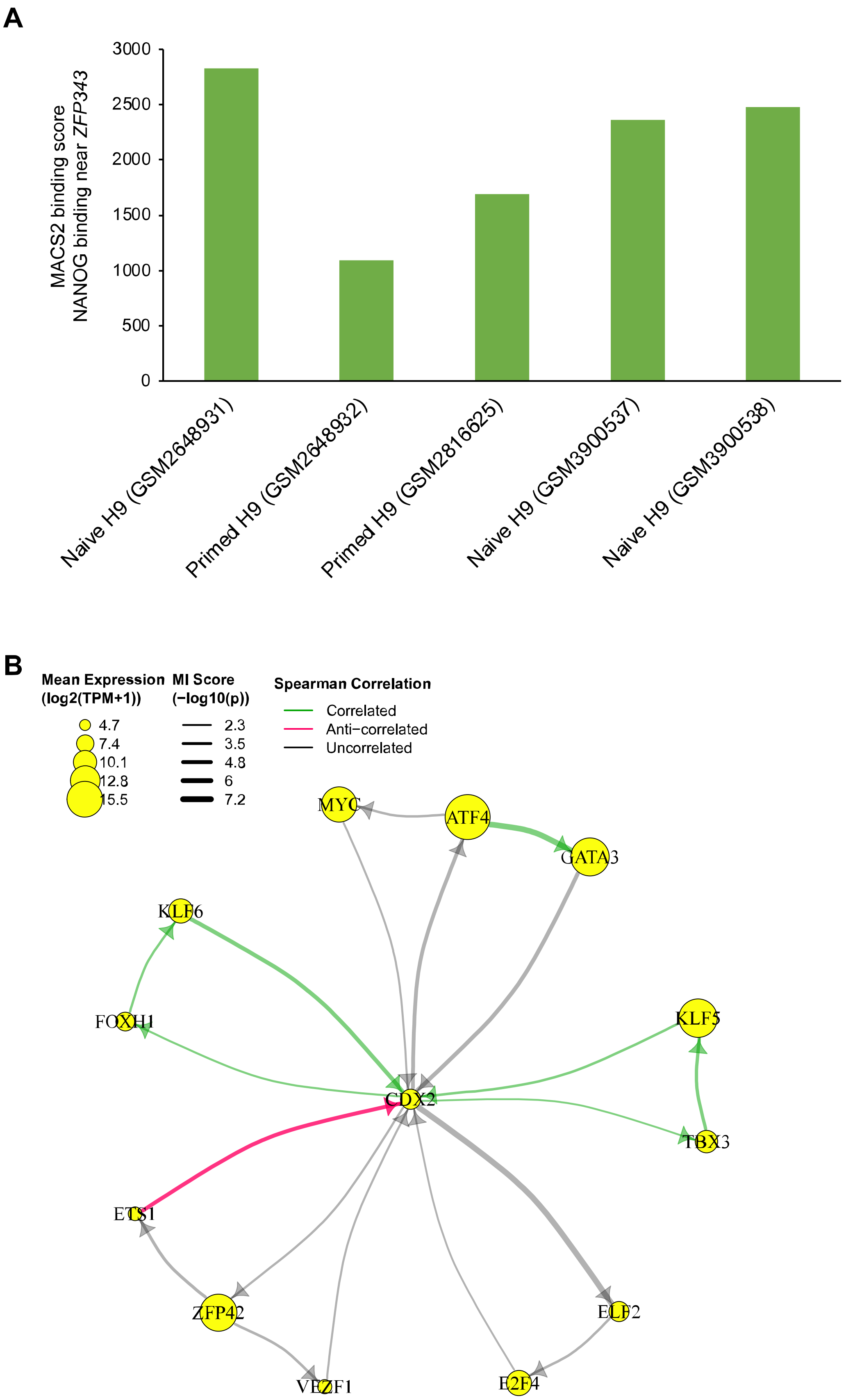


**Figure S10. (A)** MACS2 binding score to ZNF343 (±10 kb from TSS) from NANOG ChIP-seq studies in naïve and primed hESC H9 cell lines. **(B)** Feedback network formed between CDX2 targets and receptors.

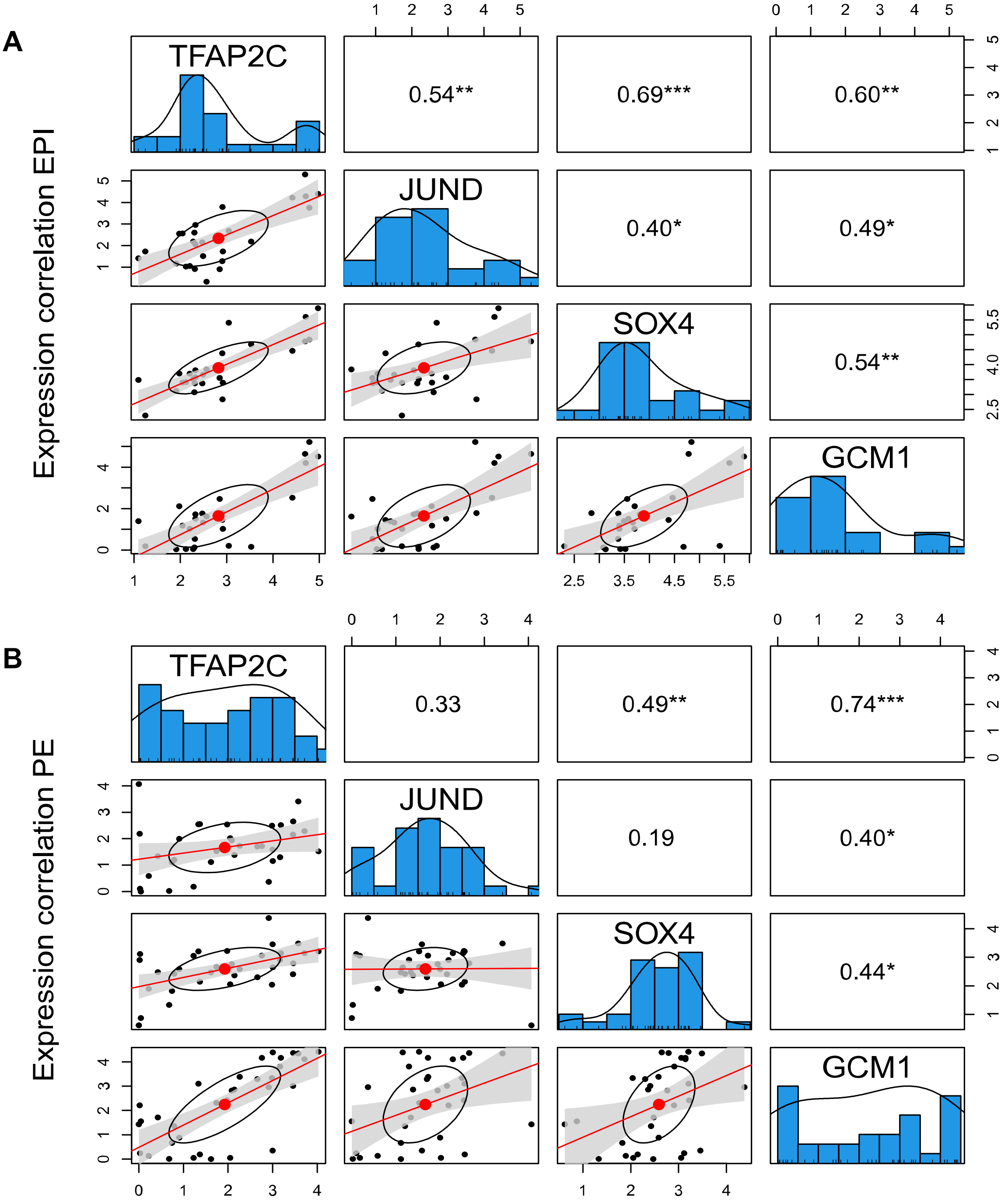


**Figure S11.** Correlation plots between TFAP2C, JUND, SOX4 and GCM1 RNA expression in the **(A)** EPI or **(B)** PE lineage of human blastocysts.

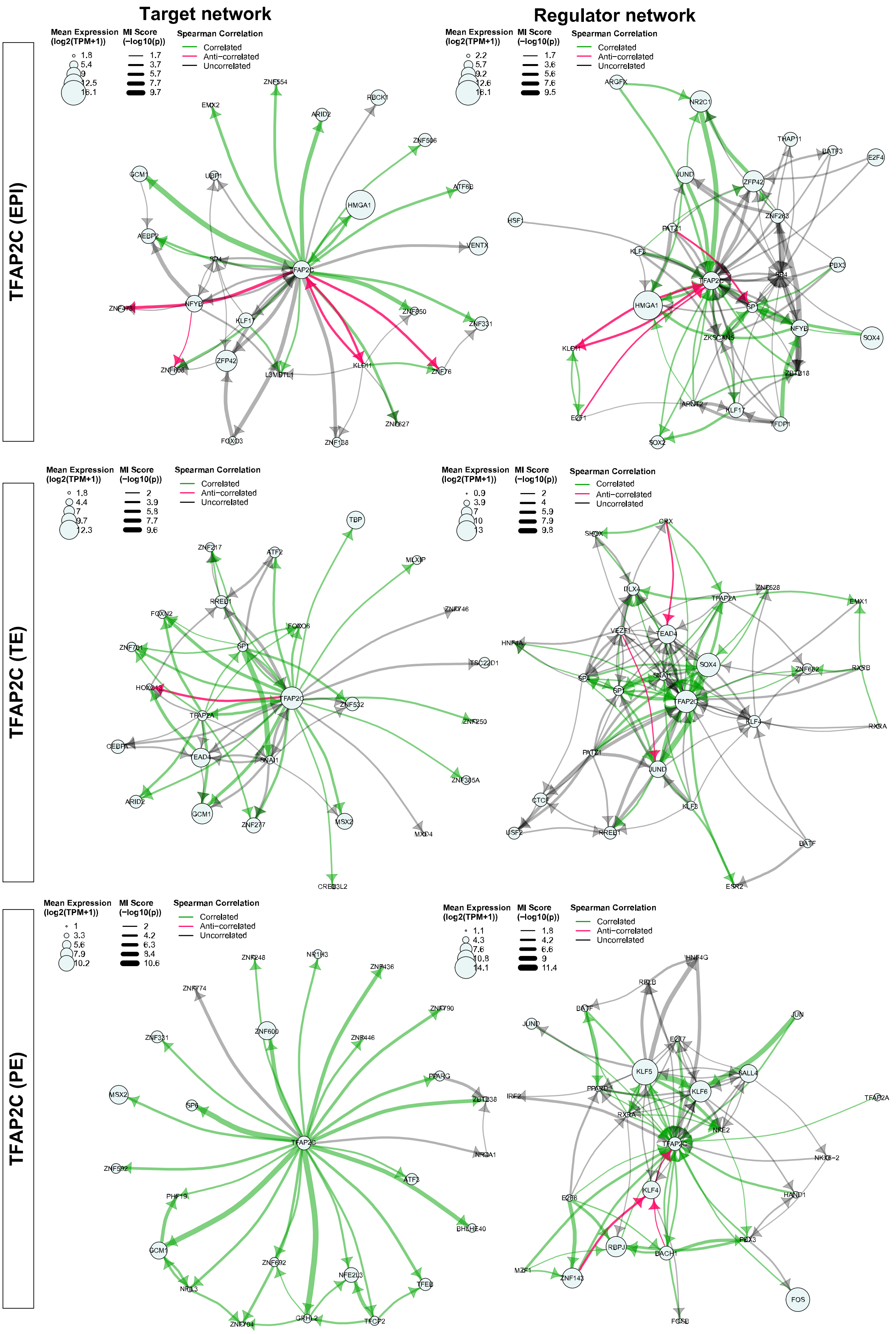


**Figure S12.** Target networks and regulator networks of TFAP2C in TE, EPI and PE cells.

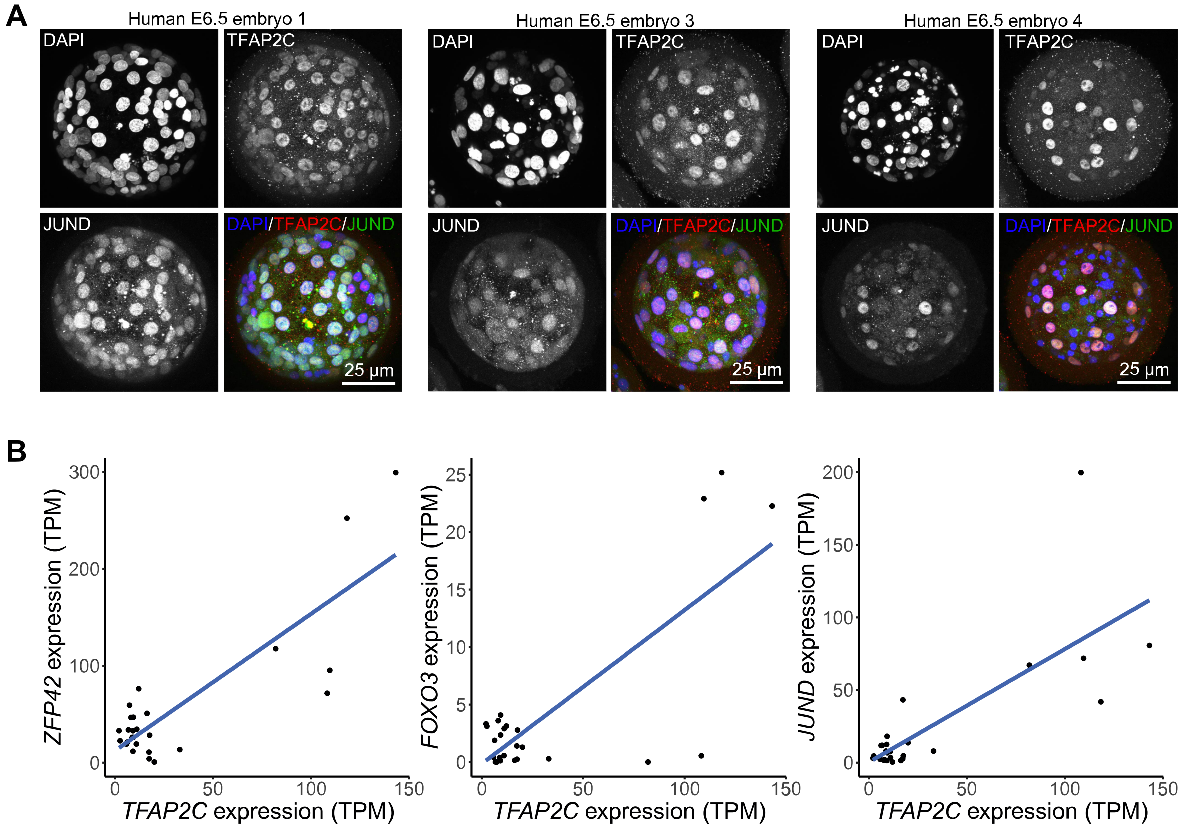


**Figure S13. (A)** Immunofluorescence staining of TFAP2C and JUND in E6.5 human blastocysts. **(B)** scRNA-seq expression plots of TFAP2C versus ZFP42 (left), FOXO3 (middle), or JUND (right).
